# Expanding kinetoplastid genome annotation through protein structure comparison

**DOI:** 10.1101/2024.08.07.607044

**Authors:** J.M. Trinidad-Barnech, J.R. José Sotelo-Silveira, D. Fernandez Do Porto, P. Smircich

**Author notes:** Correspondence: Dr. Pablo Smircich.

## Abstract

Kinetoplastids belong to the supergroup Discobids, an early divergent eukaryotic clade. Although the amount of genomic information on these parasites has grown substantially, assigning gene functions through traditional sequence-based homology methods remains challenging. Recently, significant advancements have been made in *in silico* protein structure prediction and algorithms for rapid and precise large-scale protein structure comparisons. In this work, we developed a protein structure-based homology search pipeline (ASC, Annotation by Structural Comparisons) and applied it to annotate all kinetoplastid proteins available in TriTrypDB. Our pipeline assigned functional annotation to 23,000 hypothetical proteins across all 35 kinetoplastid species in the database. Among these, we identified ubiquitous eukaryotic proteins that had not been previously detected in kinetoplastid genomes. The resulting annotations (KASC, Kinetoplastid Annotation by Structural Comparison) are openly available to the community (kasc.fcien.edu.uy).

**Author Summary:** Kinetoplastids are a group of parasites that cause severe diseases in the poorest regions of the world. Despite the increasing amount of genomic information available on these parasites, predicting the function of many of their genes using traditional methods has been difficult. Recently, there have been significant advancements in predicting protein structures and comparing them on a large scale. In this study, we created a new method called ASC (Annotation by Structural Comparisons) to find functions for all the kinetoplastid genes listed in the TriTrypDB database. Our strategy successfully assigned functions to 23,000 proteins in kinetoplastids. Among these, we discovered important proteins found in all eukaryotes that had not been previously identified in kinetoplastids. This information (KASC, Kinetoplastid Annotation by Structural Comparison) is freely available at kasc.fcien.edu.uy.

## 1 Introduction

Kinetoplastids are flagellated protozoan parasites belonging to the early branching supergroup Discobids, comprising an early divergent eukaryotic clade (Burki et al., 2020). While the amount of genomic information on these parasites has grown substantially, a significant challenge the community faces is the high percentage of genes lacking functional annotation, impeding comprehensive interpretation of genome-wide studies (Choi & El-Sayed, 2012). This deficiency in functional assignment is attributed not only to species-specific genes but also to the ancestral divergence of kinetoplastids within the eukaryotic lineage (Burki et al., 2020).

Functional annotation of proteins is crucial for understanding cellular biology at the molecular level. However, rapid increases in gene sequences driven by high-throughput technologies challenge traditional methods relying on extensive manual curation. In this context, automated annotation methods are increasingly necessary to overcome this bottleneck. Historically, functional annotation strategies depend mainly on protein sequence homology or supervised learning approaches (Position-Specific Score Matrices or Hidden Markov Models) (Altschul et al., 1997; Eddy, 2011; Remmert et al., 2012). Despite the success of sequence-based homology inference, detecting distant evolutionary relationships remains challenging for these approaches (Al-Fatlawi, Menzel, et al., 2023).

Since protein structure is more conserved than the amino acid sequence, structural alignment methods are especially relevant in phylogenetically distant organisms, offering greater sensitivity to identify homology (Illergård et al., 2009). However, the large-scale use of this approach was always limited by the availability of experimentally determined protein structure data, the difficulty of predicting protein structure from only its amino acid sequence, and the speed of structural comparison methods (Kempen et al., 2022).

Recently, significant advancements have been made in the field of *in silico* structure prediction, solving this problem for many proteins and opening new possibilities in structural biology and bioinformatics (Jumper et al., 2021; Kempen et al., 2022; Lin et al., 2023). Algorithms such as AlphaFold, RoseTTAFold, and ESMFold now allow the prediction of three-dimensional protein structures based solely on amino acid sequences, achieving resolutions competitive with experimental results (Jumper et al., 2021). The availability of comprehensive protein structures significantly enhances our ability to identify structurally similar proteins. This progress is driven by algorithms that enable rapid and precise large-scale protein structure comparisons (Kempen et al., 2022; Lin et al., 2023). These algorithms facilitate efficient protein structure searches in extensive databases like AlphaFoldDB (AFDB) (Varadi et al., 2022) and the ESM Metagenomic Atlas (Lin et al., 2023). This has allowed the development of new strategies to search for distant homology based on structural comparisons. Similar to reciprocal best hits (sequence-based, such as blast approaches), structural reciprocal best hits (SRBH) are being used to improve homology inference, functional annotation, and evolutionary analysis (Monzon et al., 2022; Svedberg et al., 2024). SRBH has shown potential in detecting novel homologs between distant model organisms within the Opisthokonta clade (Monzon et al., 2022; Svedberg et al., 2024). A comprehensive assessment of the strengths and weaknesses associated with employing these tools is ongoing. Recent analysis suggests that BLAST offers precise homolog identification with stringent E-values, hidden Markov models provide higher sensitivity but lower specificity, and structure comparison finds a balance between these extremes (Al-Fatlawi, Menzel, et al., 2023).

In this work, we propose an approach based on SRBH to annotate the genomes of highly divergent organisms from the kinetoplastid group through structural comparisons with the predicted structures from 45 model organisms. To achieve this aim, we developed a specific tool to make predictions for new genomes of kinetoplastids or other organisms, requiring little input from the user. Our results demonstrate the effectiveness of this strategy, producing concordant annotations for proteins with known functions in the database. Moreover, we assigned putative functions to thousands of genes previously classified as hypothetical in the reference kinetoplastid database. The tool (ASC, Annotation by Structural Comparison) and the resulting kinetoplastid annotation (KASC, Kinetoplastid Annotation by Structural Comparison) are openly available to the community.

## 2 Materials and Methods

### 2.1 Kinetoplastid protein sequences and structures

We manually downloaded all the protein sequences available in TriTrypDB release 65 (Shanmugasundram et al., 2023). We clustered the sequences using MMseq2 with “`foldseek cluster`” (sensitive clustering) and parameters “`--cluster-mode 1 -- similarity-type 2 --min-seq-id 0.5 -c 0.8 --cov-mode 0 -e 1e-5`”, as in Barrio-Hernandez et al. 2023. We considered all protein sequences in each cluster as homologs. Clusters with less than ten protein sequences were discarded. All available structures for each sequence in the remaining clusters were obtained via FTP from the AFDB. Since clusters can contain multiple structures, we calculated the average pLDDT for each structure and selected the one with the highest as the cluster representative (using in-house Python scripts available on https://github.com/JuanTrinidad/ASC), as in Barrio-Hernandez et al. 2023.

As inconsistencies between the sequences from TriTrypDB and AFDB were observed for some proteins, structure files were curated by comparing their sequence to the one from TriTrypDB. If the length of the PDB and the TriTrypDB sequences were identical, they were deemed matching. Conversely, both protein sequences were aligned if a length discrepancy was observed. Sequences exhibiting less than 80% identity were excluded from consideration.

### 2.2 Proteins structure data from model organisms

The protein structures of model organisms used in this work were downloaded from “AFDB model organisms proteomes” (https://ftp.ebi.ac.uk/pub/databases/alphafold/v4/). We excluded kinetoplastid proteomes, obtaining a final number of 45 proteomes (Supplementary Table 1).

### 2.3 Protein structure comparison

Protein structures were compared using Foldseek release 8.ef4e960 (Kempen et al., 2022). First, query (14,267 protein structures representative of all kinetoplastid clusters) and target (one for each independent model organism) databases were created using the “`foldseek createdb`” command. Reciprocal best hit (RBH) comparisons were performed independently between the query database and each target database using “`foldseek rbh`” with parameters “`-s 9.5 -c 0 -a`”.

Structural alignment was performed with the FATCAT algorithm with default parameters (Flexible structure AlignmenT by Chaining Aligned fragment pairs, allowing Twists) (Li et al., 2020), and the TM-score was calculated using the TM-align algorithm (Zhang & Skolnick, 2005). Alignments were visualized using ChimeraX software (Pettersen et al., 2004).

### 2.4 Functional annotation validation

To validate our results, for all kinetoplastid cluster representatives and all model organism proteins, we downloaded from UniProt the protein functional annotation of the Protein Families, PANTHER, InterPro, and Pfam databases (The UniProt Consortium, 2023) (UniProt: the Universal Protein Knowledgebase in 2023). For each SRBH, when available, we compared query and target functional annotation by creating five categories. Sets are defined as Q (set of functional annotations for the query SRBH protein) and M (set of functional annotations for the model organism protein).

Categories:

- ALL: Q = M
- K_in_M: Q ⊆ M
- M_in_K: M ⊆ Q
- PARTIAL: (Q ∩ M) ≠ Ø and Q ≠ M and M ≠ Q
- ZERO: Q ∩ M = Ø

Gene Ontology enrichment analysis was performed and visualized by WEGO 2.0 (Web Gene Ontology Annotation Plot) (Ye et al., 2018).

### 2.5 Validation of protein subcellular localization prediction using TrypTag

Our results were also evaluated using the TrypTag database, which has experimental protein targeting and localization data for almost all proteins coding genes in *T. brucei*.

We downloaded the subcellular localization data (Gene Ontology, cellular component) from UniProt and the experimental data from TrypTag from TryTripDB (Billington et al., 2023). We selected nine ubiquitous subcellular localizations for the comparison: cytoplasm, nucleoplasm, nucleolus, basal body, endoplasmic reticulum, mitochondrion, nuclear envelope, nucleus, and Golgi apparatus. We compared the semantic similarity between GO terms from UniProt and TrypTag using GOGO (Zhao & Wang, 2018). Before the comparison, GO terms from TrypTag with percentages or weak annotations “weak|10%|25%|50%|75%” were considered low-confidence annotations and removed. Given that many genes possess multiple ontology terms within the UniProt annotation (frequently nested), we conducted a comprehensive comparison against the TrypTag annotation. For each gene, using GOGO, each Uniprot GO term obtained by structural annotation was compared to all GO terms annotated in TrypTag. We defined the GOGO score between both annotations as the highest score obtained in each comparison.

The comparisons of score distributions from TrypTag validation were tested using the Mann-Whitney-U tests from the SciPy library (Jones Eric & Pearu Peterson, 2001) and visualized with the Seaborn library.

### 2.6 BUSCO analysis

We used the Benchmarking Universal Single-Copy Orthologs algorithm (BUSCO) (Manni et al., 2021)to identify essential eukaryotic genes reported as missing in kinetoplastids. For this, we ran BUSCO on the TriTrypDB protein fasta file using the “`-m prot`” option and the `eukaryota_odb10` database. For comparison, we performed the same procedure for model organism proteomes.

## 3 Results

### 3.1 ASC (Annotation by Structural Comparisons) Pipeline Description

The entire workflow was implemented in the Snakemake workflow management system (Köster et al., 2021), a popular tool for automating bioinformatics workflows. The execution script and the required dependencies are available at https://github.com/JuanTrinidad/ASC. A configuration file located in the ‘config’ directory allows for the adjustment of all parameters mentioned in the methods section (as well as others configurable according to the documentation of each software). This pipeline was developed not only to run the analyses presented in this manuscript but also to provide a tool that users can easily implement for their organism of interest.

The workflow requires two files: a fasta file containing the protein sequences to be annotated and a .tsv file linking the fasta headers with a UniProt accession. The workflow initiates by clustering the protein sequences to be annotated using MMseq2 (Steinegger & Söding, 2017) (see Materials and Methods) and setting the minimum number of proteins per cluster to be retained. Once the sequences are clustered, the clusters are filtered by the number of sequences defined by the user. Using the second mandatory TSV file, the pipeline downloads all available structures for the sequences within the clusters via FTP from AFDB. Each protein structure will be compared with the sequences in the fasta file to prevent inconsistencies that might exist between sequences in cases where sequences are obtained from different databases (see Materials and Methods). Since clusters can contain multiple structures, we choose the best structure prediction for each cluster based on the average pLDDT. The pipeline then creates a database for the Foldseek program using these representative structures (query database). The remaining clusters without available protein structures are excluded from the analysis.

In parallel, the Snakemake script downloads all available model organisms from AFDB and creates an independent database for each organism. These databases are used as targets. Using foldseek reciprocal-best-hit (equivalent to SRBH), comparisons are performed independently between the query database and each target database. For each query, the pipeline selects the top number of SRBH as specified by the user (sorted by Foldseek e-value) and performs a flexible structural alignment between the corresponding PDB files using the FATCAT algorithm (Li et al., 2020). FATCAT is a structure aligner that allows for flexibility, an important aspect because AlphaFold structures often have different domain topologies/orientations (Xia et al., 2023). The TM-score is then calculated for these aligned structures using the TM-align algorithm (Zhang & Skolnick, 2005) (Figure 1). The final result consists of a table with all SRBH results from FoldSeek and a separate table with the results from TM-align and all structural comparisons made with FATCAT as separate PDB files.

**Figure 1.**
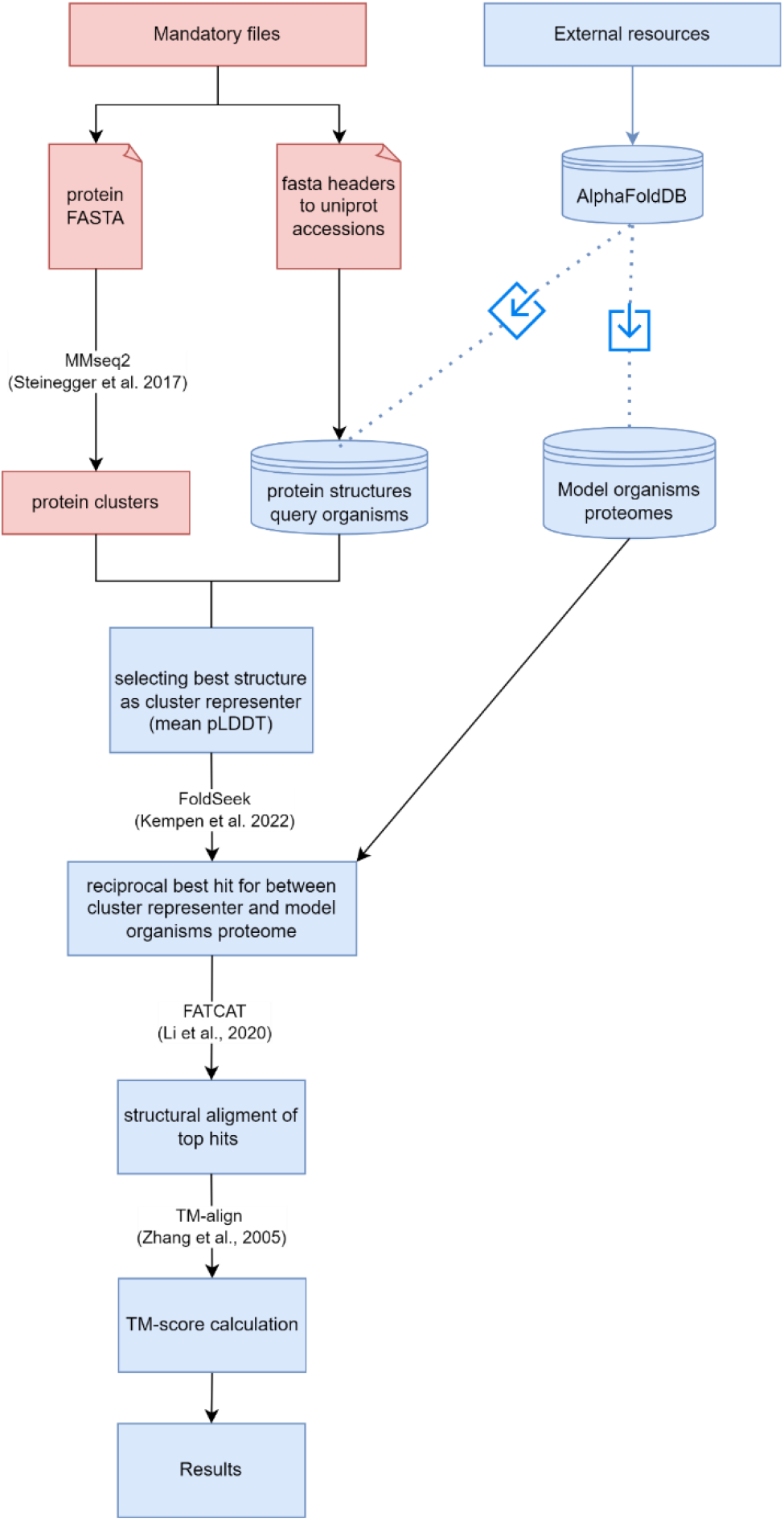
ASC pipeline workflow. Workflow outlining the process for functionally annotation of proteins using structural data. The pipeline begins with mandatory files, including protein sequences (as FASTA) and the corresponding UniProt accessions. MMseq2 is employed to generate protein clusters. All protein sequences with available protein structures in AFDB will be downloaded, and the best structural representative for each cluster will be selected based on mean pLDDT scores. FoldSeek is used to identify structural reciprocal best hits between cluster representatives and model organism proteomes. Top hits undergo structural alignment using FATCAT, followed by TM-align to calculate TM-scores, and filtering leading to the final results.

### 3.2 Structural annotation of kinetoplastid genomes

The development of this pipeline enabled us to analyze 657,192 kinetoplastid protein sequences from all 84 genomes available in TriTrypDB. Proteins were clustered, and each group was filtered by size, with a cluster considered valid if it contained at least ten sequences. This criterion was used to avoid further analysis of incorrect gene predictions or genes lacking homology across genomes. These initial clustering and selection steps discarded 117,582 hypothetical proteins and 61,198 annotated proteins, potentially representing species-specific genes with insufficient genomes to meet the minimum number of homologs, pseudogenes, highly variable sequences, or artifacts generated during genome annotation. This process resulted in 14,778 clusters encompassing 478,612 sequences, of which 210,201 (44%) were hypothetical proteins. Among these, 3,565 clusters, comprising 86,679 sequences, consisted solely of hypothetical proteins (termed “dark clusters” as defined by Barrio-Hernandez et al. 2023). The remaining 123,522 hypothetical proteins were grouped with previously annotated protein sequences, representing 59% of the hypothetical proteins. So, due to our stringent clustering parameters we were able to assign annotations to a significant number of hypothetical proteins based solely on sequence homology (Barrio-Hernandez et al., 2023; Pearson, 2013).

From the 14,778 retained clusters, 14,267 had protein structures available in AFDB, which served as our query database for subsequent pipeline steps. The remaining 511 clusters, lacking available protein structures, were excluded from the analysis. AFDB contains only a few kinetoplastid genomes, so these clusters likely lacked genes from them, resulting in their exclusion from assigned protein structures. Currently, we are in the process of modeling these proteins to incorporate them into future versions of the database. For the clusters where a representative structure could be obtained, we performed the SRBH step against model organisms retaining 11,753 protein clusters, including 2,689 dark clusters. We selected the top five SRBH for each query, conducted structural alignments using FATCAT against the top targets reported by Foldseek, and calculated the TM-score. Using a stringent TM-score threshold of 0.5 for protein homology (Xu & Zhang, 2010), we filtered the results (both TM-score Chain 1 and TM-score Chain 2 > 0.5), creating our “Final Dataset.” It is worth noting that the TM score considers the alignment length between hits. Our chosen parameters prioritize full protein alignments, excluding partial alignments that might arise from similar subdomains. The final dataset, derived from our rigorous pipeline steps and TM-score filtering, resulted in 7,486 clusters, including 942 dark clusters comprising 23,290 protein sequences (Figure 2). These clusters are particularly interesting because they appear in multiple genomes or copies within a single genome and lack homology with genes annotated in TriTrypDB. Notable examples are discussed in the last section of the article.

**Figure 2.**
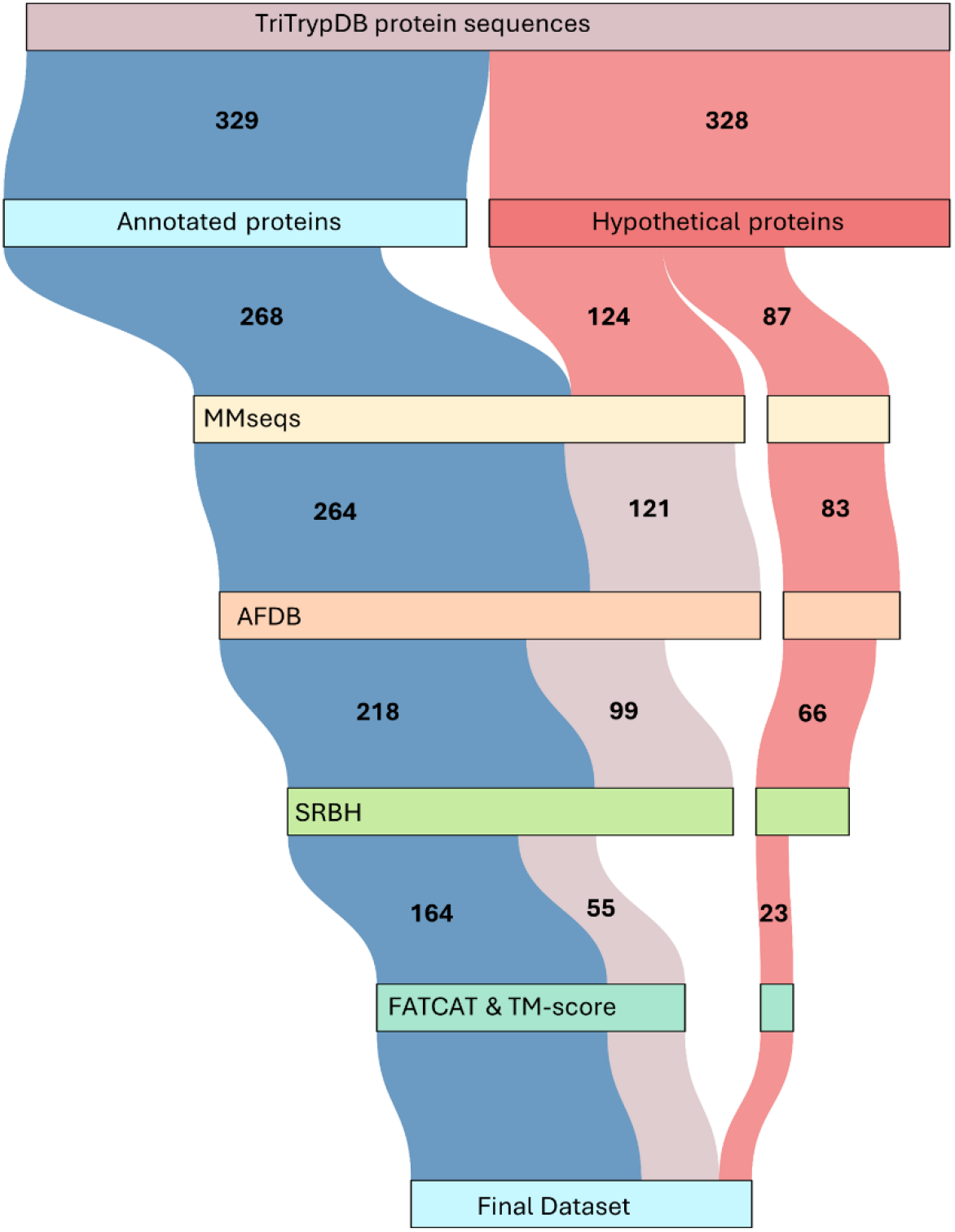
The number of protein sequences retained throughout the main steps of the pipeline. Sankey diagram visualizing the number of genes retained after each main step of the functional annotation pipeline. The nodes in the diagram, each represented by a specific color, denote the pipeline’s stages, while the links between nodes indicate the number of genes (in thousands) transitioning from one step to the next. The color coding distinguishes between hypothetical and annotated genes as per TriTrypDB: red represents hypothetical genes, blue represents annotated genes, and light red represents hypothetical genes with sequence homology to annotated genes. The process begins with 329K annotated proteins and 328K hypothetical proteins from TriTrypDB. After clustering with MMseqs2 and selecting clusters with more than 10 members, 268K annotated proteins and 124K hypothetical proteins share clusters, while 87K hypothetical proteins remain in Dark Clusters. Following clustering, protein structures are retrieved from AFDB. At this stage, 264K annotated and 121K hypothetical proteins from the clustered sets advance, along with 83K hypothetical proteins from Dark Clusters, indicating the availability of a structure for the cluster representative. The workflow then identifies SRBH, where 218K annotated proteins and 99K hypothetical proteins meet the criteria, with an additional 66K hypothetical proteins retained from Dark Clusters. The final step involves performing structural alignment using FATCAT and calculating TM-scores to filter the SRBH. The workflow effectively filters and refines the initial protein sequences, resulting in a robust final dataset comprising 164K annotated plus 55K hypothetical proteins, and 23K hypothetical proteins derived from Dark Clusters.

As expected, GO analysis of the terms associated with these new annotations revealed a diverse range of functional categories. Notably, several categories such as membrane-associated were overrepresented compared to their frequency in the previously annotated terms (Supplementary Figure 1).

It is important to emphasize that, irrespective of whether a function can be assigned to these clusters based on the model organism’s annotation, finding homology suggests that these genes are conserved among highly divergent organisms and thus likely have fundamental or essential functions.

The results are accessible at kasc.fcien.edu.uy. The default tab allows users to query a GeneID, displaying TritrypDB and UniProt annotations, the top five structural comparison results, and a visualization of the structural alignment of the queried protein against a selected organism. The secondary tab provides information on the corresponding cluster that the ID belongs to and a detailed description of the structural alignment.

### 3.3 Validation of functional annotation

To validate our approach, we used the Protein Families, PANTHER, InterPro, and Pfam databases as references. We assessed the overlap of annotation of all SRBH with functional annotation information in the different databases (see Methods).

Our results from the Protein Families and PANTHER databases show that our annotation is highly consistent with previous protein family-level annotations. The data suggest an enhancement in annotation precision after filtering, as seen by the increased number of exact matches between annotations (ALL category) and the reduction of non-overlapping annotations considered incorrect assignments (ZERO category) (Figure 3 panels A and B). In the first case, we obtain a 90% success rate before filtering, increasing to 95% when we perform filtering by TM-score. These results are similar to those of the PANTHER database at the superfamily level where we are at 77% accuracy without filtering, increasing to 88% in the Final Dataset (Figure 3 panels A and B). Therefore, we are facing a reliable family prediction that is comparable to the results obtained by previous studies in organisms even more closely related to each other (inside the Opisthokonta clade) (Monzon et al., 2022).

**Figure 3.**
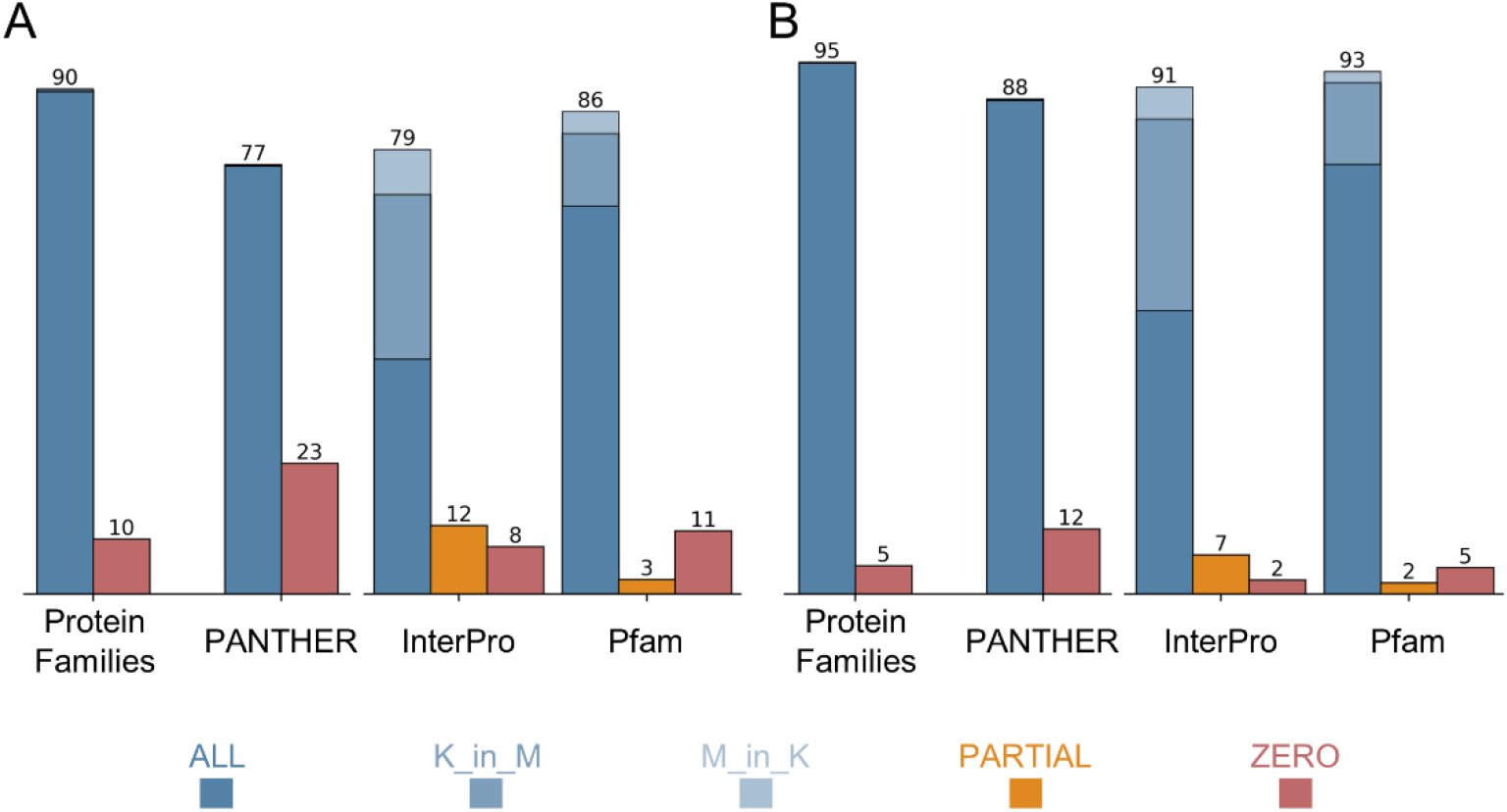
Evaluation of SRBH annotations before and after filtering. Comparison of reciprocal best hit (RBH) annotation results for query and target proteins using four databases: Protein Families, PANTHER, InterPro, and Pfam. The left panel (A) displays the annotation results before applying a filtering process, while the right panel (B) shows the results after filtering (Final Dataset). Each bar represents the percentage of annotations categorized as ALL (blue), K_in_M (light blue), M_in_K (lighter blue), PARTIAL (orange), and ZERO (red). The numbers at the top of each bar indicate the total percentage of the column rounded.

At the domain level, we also have an increase in the ALL and a decrease in the ZERO and PARTIAL categories. Using the InterPro and Pfam databases, within the ALL category, we distinguish between exact matches (dark blue) and matches where the domains of one protein are a subset of the other (light blues, see Methods). As shown in Figure 3, we observe 79% and 86% coincidence without filtering, rising with the Final Dataset to 91% and 93%.

It is worth noting that the light blue data is mostly composed of cases where the query protein annotation is a subset of the target one (K_in_M). The lack of annotation in kinetoplastids explains the K_in_M category percentage, so our pipeline results in a considerable advance for proteins without any annotation (see last section) and genes that already have domains annotated.

Considering this in our Final Dataset, we are deepening the annotation of 34% of the clusters in InterPro and 15% in Pfam (K_in_M). In the opposite case, where an annotation in model organisms is not observed in the kinetoplastid protein, it is more difficult to affirm that this is due to a lack of annotation. However, the percentage of cases is low. The PARTIAL and ZERO matches are low, as both added do not reach 10% if we consider after filtering.

### 3.4 TrypTag validation

TrypTag is a genome-wide experimental database that details protein locations within the *T. brucei* parasite. Researchers used fluorescent tags and microscopy to determine protein locations, and each cell line was manually annotated using a standardized ontology (Billington et al., 2023). This database is a highly accurate reference for our approach, enabling us to validate our tentative annotations against a gold standard.

To enhance the reliability of our analysis, we removed non-confident entries and selected ubiquitous GO cellular components (CC) from TrypTag (see Methods) and used the GO CC annotation from our Final Dataset to compare. The results from the SRBH analysis can exhibit various combinations of annotations in the GO cellular component category: only the kinetoplastid protein, only the model organism protein, both, or neither may be annotated. Whenever possible, we independently compared the annotations of the kinetoplastid and model organism proteins obtained from UniProt with those assigned by TrypTag for each SRBH. We assessed the semantic similarity of the GO terms using GOGO (Zhao & Wang, 2018). Since the model organism’s annotation theoretically represents the final annotation result for our strategy, we used this comparison to evaluate the accuracy of our approach. In other words, we evaluated whether our annotations are comparable to the existing annotations for kinetoplastid proteins.

Figure 4 shows the distribution of semantic similarity values of GO terms for each comparison. The grey violin plots represent the distribution of GOGO scores by comparing the annotation of kinetoplastid genes (previous annotation) with the annotations assigned by TrypTag, which serves as our reference. The red violin plots represent the results from comparing the annotation data of model organisms with the annotations of TrypTag, which approximates the confidence level we can obtain with our annotation strategy for hypothetical gene clusters.

**Fig. 4.**
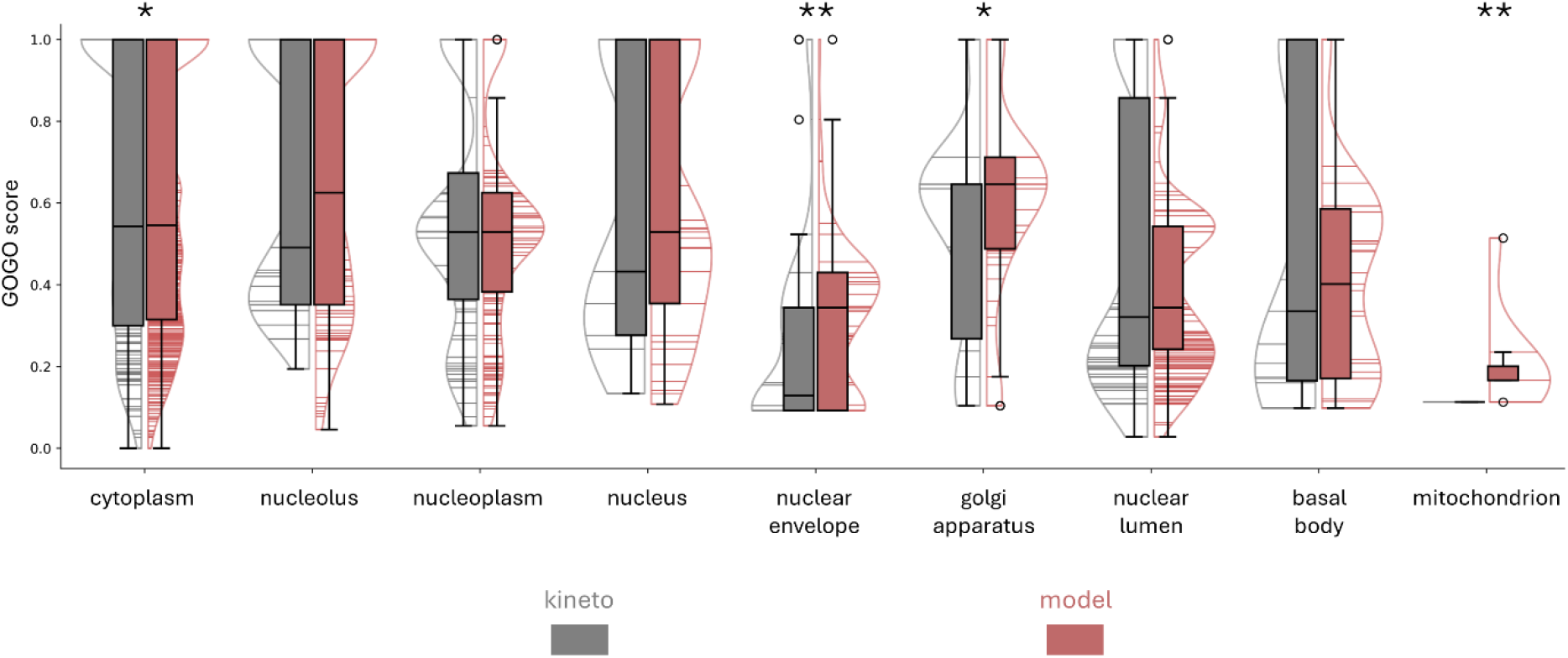
Comparison of GO cellular component semantic similarity scores for kinetoplastid and model organism annotations. Distribution of GO term semantic similarity scores (GOGO scores) for two categories: kinetoplastid (grey) and model organism (red). Each violin plot displays the range and distribution of scores for ubiquitous cellular components, including cytoplasm, nucleus, nuclear lumen, nuclear envelope, nucleolus, basal body, golgi apparatus, and mitochondrion. The boxplot within each violin plot represents the interquartile range (IQR) and the white lines indicate the median values. * p-value < 0.05, ** p-value < 0.01

Using the Mann-Whitney-U test to compare the means, we did not observe significant differences for most categories, meaning that the current annotation is equally coherent with TrypTag as our analysis. We observe a trend for cytoplasm, nuclear envelope, and Golgi apparatus, indicating that SRBH strategy annotates subcellular localizations more in line with the TrypTag experimental results.

In conclusion, our annotation is at least as good as the currently accepted methods when compared against TrypTag, and it contributes new information for previously uncharacterized proteins.

### 3.5 The missing gears

After validating our approach, we sought biologically relevant examples within our results, focusing on Dark Clusters and Annotated Clusters lacking precise annotations. To refine our search and identify truly hypothetical clusters using various databases, we incorporated functional annotation data from UniProt, specifically the Product Description, in addition to TriTrypDB.

We ran the BUSCO eukaryotic database (Manni et al., 2021) for all available protein sequences in TriTrypDB. This database contains 255 “BUSCO groups,” of which 68 were reported as “Missing.” Our structure comparison annotation approach identified 48 out of these 68 missing BUSCO groups. Notably, many of these genes were annotated in TriTrypDB or UniProt, likely through other annotation methods such as experimental evidence or BUSCO databases. However, by manually inspecting BUSCO results of model organisms and their SRBHs, we identified 10 BUSCO groups not found by any other functional annotation methods in kinetoplastids; examples are provided in Figures 5 and Supplementary Figure 2. Gene IDs for all proteins belonging to these clusters are provided in Supplementary Tables 2 and 3. These genes span distinct core functional categories, including transcription and DNA repair, translation, and cell cycle regulation. In transcription and DNA repair categories, the General Transcription and DNA Repair Factor IIH subunit TFB4 is a crucial component of the TFIIH core complex involved in DNA damage response, repair, and transcription regulation (Rimel & Taatjes, 2018). The RNA Polymerase II subunit A, previously reported as absent in kinetoplastid (Pons et al., 2014), acts as a protein phosphatase and is critical in mRNA processing and termination (Cossa et al., 2021). Additionally, the Cleavage Stimulation Factor subunit 3 is required for polyadenylation and 3’-end cleavage of mammalian pre-mRNAs (Takagaki & Manley, 1994, 2000). Considering translation related functions, the Translational Activator GCN1 functions as a ribosome collision sensor, activating translation quality control and the integrated stress response (Oltion et al., 2023), while the tRNA (guanine-N(7)-)-methyltransferase non-catalytic subunit plays a role in tRNA and mRNA processing (Jin et al., 2023). For cell cycle regulation, we found the E3 Ubiquitin-Protein Transferase MAEA, a core component of the CTLH E3 ubiquitin-protein ligase complex, which is crucial for ubiquitination, proteasomal degradation, and cell proliferation (Lampert et al., 2018). Lastly, in protein assembly and mitochondrial functions, the Ubiquinol-Cytochrome C Chaperone is vital for the assembly of Ubiquinol-cytochrome C Reductase in the mitochondrial respiratory chain (Gruschke et al., 2012), and the Iron-Sulfur Cluster Co-Chaperone Protein HscB is involved in iron-sulfur cluster assembly and protein maturation (Maio et al., 2016; Uhrigshardt et al., 2010), contributing to mitochondrial electron transport chain function.

**Figure 5.**
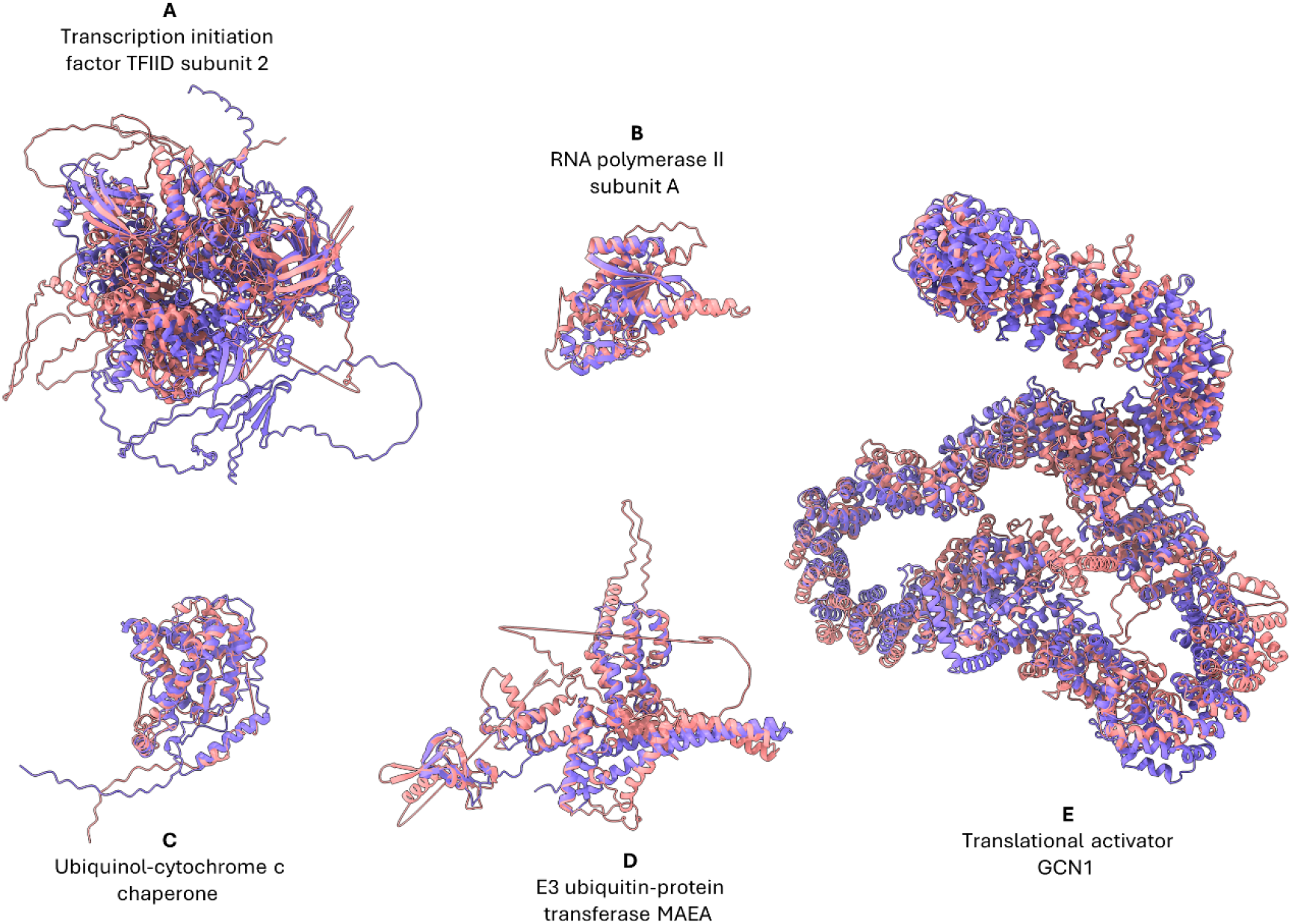
Essential proteins for eukaryotes discovered through our approach. The target protein (model species) is shown in purple, and the query protein (kinetoplastids) is shown in pink. In most cases, there are SRBH with homologs of this protein in different organisms; we chose one as an example. (A) Transcription initiation factor TFIID subunit 2, A0A1D8PQF6_CANAL vs E9AGI7_LEIIN. (B) RNA polymerase II subunit A, Q9VWE4_DROME vs Q4DKS2_TRYCC. (C) Ubiquinol-cytochrome c chaperone (CBP3), P21560 CBP3_YEAST vs A4I4I7_LEIIN. (D) E3 ubiquitin-protein transferase MAEA, Q7L5Y9 MAEA_HUMAN vs Q4D4T7_TRYCC. (E) Translational activator Gcn1, Q54WR2 GCN1_DICDI vs A4HXV6_LEIIN

Of these 10 BUSCO groups, 5 had SRBHs with a single kinetoplastid cluster within our database (Figure 5), while 5 had multiple SRBHs (Supplementary Figure 2). Examining the four sequence clusters with multiple SRBHs, we observe that the clusters are grouped by genus, with Trypanosoma generally being distinct from the other genera (Supplementary Table 4). This is primarily due to the stringent criteria regarding sequence identity and coverage used for clustering kinetoplastid proteins, often resulting in the division of homologous genes into subclusters. If only one target model species were used, only one of the clusters would represent the SRBH. However, by using several model species, we obtained multiple SRBHs for the same BUSCO group or gene homologs. None of the structural alignments in Figures 5 and Supplementary Figure 2 exceeded a sequence identity of 20% (except for some hits from co-chaperone Hsc20), demonstrating the high level of divergence and, consequently, the difficulty in establishing homology relationships through classical methods (see Supplementary Table 4). The subcellular locations, according to TrypTag, of the missing genes are shown in Supplementary Figure 3.

## 4 Discussion

The field of genomic functional annotation is undergoing a paradigm shift, partly due to the emergence of methods that reveal remote homology between genes (Al-Fatlawi, Menzel, et al., 2023). Specifically, the advent of algorithms for predicting tertiary structure and conducting efficient structural comparisons has introduced a novel approach to assessing protein similarity. It has been demonstrated that sequences-dependent bioinformatics methods cannot functionally annotate essential eukaryotic proteins in cases of high degree of sequence divergence (Al-Fatlawi, Schroeder, et al., 2023; Moi et al., 2022; Wan et al., 2017). Therefore, developing new methods based on structural comparison known to overcome these limitations holds great potential for assigning putative functions to many currently unannotated genes through homology inference to highly divergent genes (Al-Fatlawi, Menzel, et al., 2023; Borujeni & Salavati, 2024; Monzon et al., 2022; Svedberg et al., 2024). While the strengths and limitations of these methods are currently being evaluated (Al-Fatlawi, Menzel, et al., 2023; Monzon et al., 2022; Svedberg et al., 2024), they have proven to be valuable tools for elucidating the functions of previously uncharacterized proteins. Kinetoplastids offer a unique opportunity for these approaches due to their early divergence in the eukaryotic tree, the significant divergence between groups (e.g., Leishmania and Trypanosoma), and the limited scope of current functional annotation.

The bioinformatics pipeline described here (ASC) aims to facilitate structural inference of homology, allowing the annotation transfer from sequence divergent homologous proteins. ASC relies on SRBH instead of all best hits creating a stringent criterion for identifying homologous proteins and increasing precision (Monzon et al., 2022). As this results in lower sensitivity there is still room to delve deeper into the annotation by compromising some of the specificity given by the SRBH strategy. The strategy requires a database of protein structures from model species making the selection crucial; in this work we used species representative of diverse evolutionary lineages.

To increase computational efficiency, we select a representative homolog from highly similar genes for the subsequent structural homology search. The high level of divergence within kinetoplastids explains that during the initial clustering step, we noticed the sub-clustering of homologous genes within kinetoplastids. The comparison of each representative to each model organism separately can compensate for some of the issues raised by this in the context of RBH strategy. Lowering the sequence identity threshold to cluster more dissimilar sequences could partially solve this issue but this would come with an increase in false positive homology annotation (20-35% sequence identity) (Rost, 1999) and was not deemed appropriate. Although adding a structure-level clustering step after sequence-level clustering could also address this, it would be computationally expensive, even though this can be implemented in future pipeline versions.

An intrinsic limitation of our approach relies on the fact that kinetoplastids are not part of the model species set, so our approach will not yield results for kinetoplastid-specific proteins, such as those coding for surface protein families. An interesting perspective related to performing structure-based comparisons within kinetoplastids, is that the identification of homologs and, eventually, orthologs, will enhance future phylogenetic studies and comparative genomics among this group. This is particularly true for identifying genes with greater divergence or rate of evolution and the processes underlying them.

In the current work, we applied the ASC pipeline to all available kinetoplastid genomes. Considering the hypothetical proteins that passed the clustering and cluster selection steps, our work was able to add functional annotation to most of them. Among these, many were part of clusters where at least one member already had prior functional annotations. Therefore, for this group, our pipeline has effectively increased the functional annotation of current hypothetical proteins by transferring information from homologous genes determined by sequence similarity within kinetoplastid genomes. The remaining comprised hypothetical proteins that were found in Dark Clusters that matched known structures in model organisms. This group totals around 23,000 kinetoplastid proteins from 942 clusters. We consider this number to be significant, underscoring the relevance of the structural homology search approach for the functional annotation of kinetoplastid protein-coding genes. Even when the functional annotation of the model species protein hit is limited, a match suggests that the kinetoplastid hypothetical protein is not an annotation error and points to the protein’s relevance to the parasite, given its evolutionary conservation.

It is worth noting that validating *in silico* genome annotation of dark matter proteins presents significant challenges due to the absence of a broad and definitive gold standard for comparison (Ardern et al., 2023). Our strategy employed two alternative approaches. First, we analyzed currently annotated kinetoplastid proteins. This shows that the annotation obtained by our approach aligns well with previous annotations. Furthermore, our results reveal that our strategy often provides a deeper functional understanding by assigning new InterPro and Pfam domains to currently annotated genes. This additional information enhances our knowledge beyond what databases currently provide. Second, we leveraged TrypTag, a genome-wide experimental database of protein subcellular locations, achieving similar results. This demonstrates that our approach yielded comparable or superior results to existing annotation methods. Remarkably, our method was capable of transferring annotation from model species to a significant number of previously unannotated proteins present in dark clusters of kinetoplastid genomes.

Building on this new annotation, we investigated whether essential eukaryotic proteins currently absent in kinetoplastid genome annotation could be identified. Our hypothesis was that the absence of these functions might be due to sequence divergence rather than an inherent biological peculiarity. To address this, we employed BUSCO to identify essential eukaryotic genes not detected in kinetoplastid genomes. The combination of the tool with our results allowed us to identify ten groups with functions crucial for eukaryotic biology, all present in Dark Clusters. These groups encompass a range of essential processes, including transcription, translation, DNA repair, and cell cycle regulation. These results further support the significance of employing structural comparisons in divergent organisms.

## 5 Acknowledgements

We would like to thank Dr. Laurie Read for her helpful suggestions.

## 6 Author contributions

JTB, PS: Design of the methodology

JTB: Performed the analysis, developed web tool

JSS, DFD, JTB, PS: Wrote and reviewed the manuscript

JSS, PS: Acquisition of financial support

PS: Coordinated the project

