## Supplementary Figures for "Expanding kinetoplastid genome annotation through protein structure comparison"

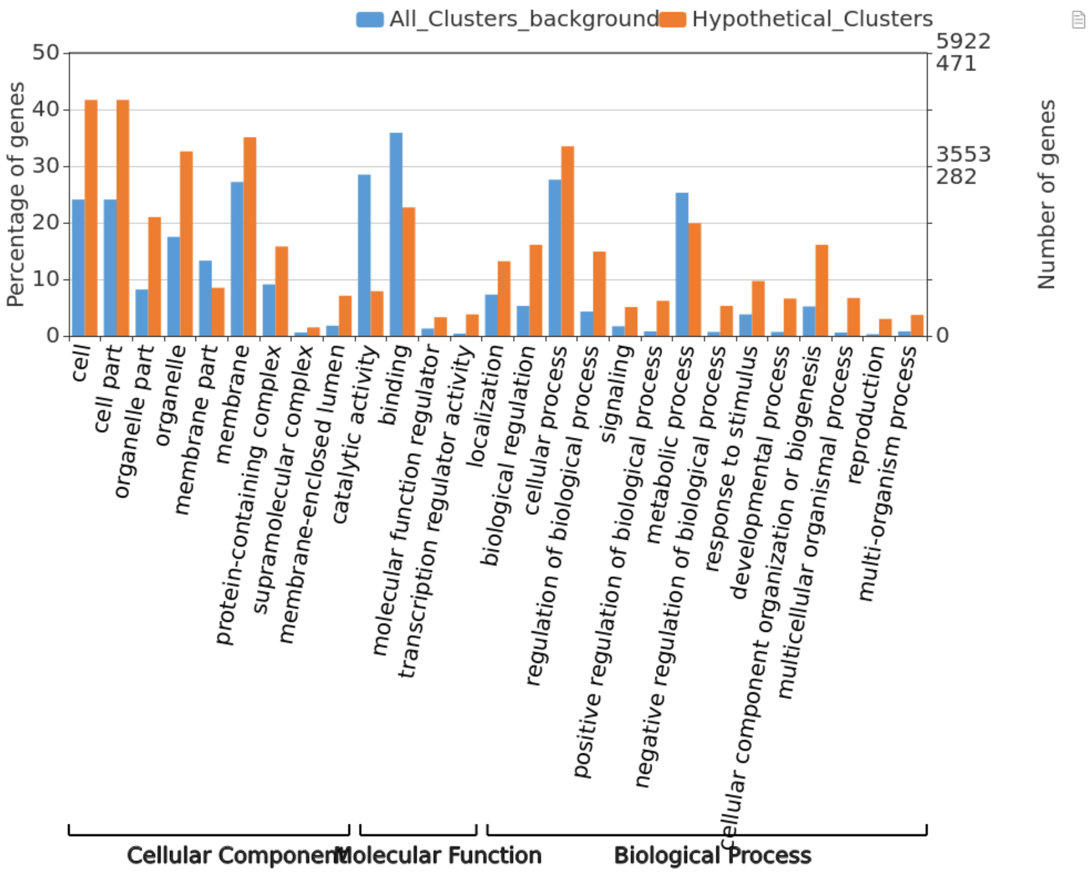


**Supplementary Figure 1. Gene Ontology terms enrichment analysis in the 942 hypothetical cluster.** This supplementary figure shows a gene ontology term enrichment analysis performed with WEGO2.0 comparing the 942 hypothetical protein clusters annotated by our method against all clusters. The analysis includes biological processes, molecular functions, and cellular components categories.

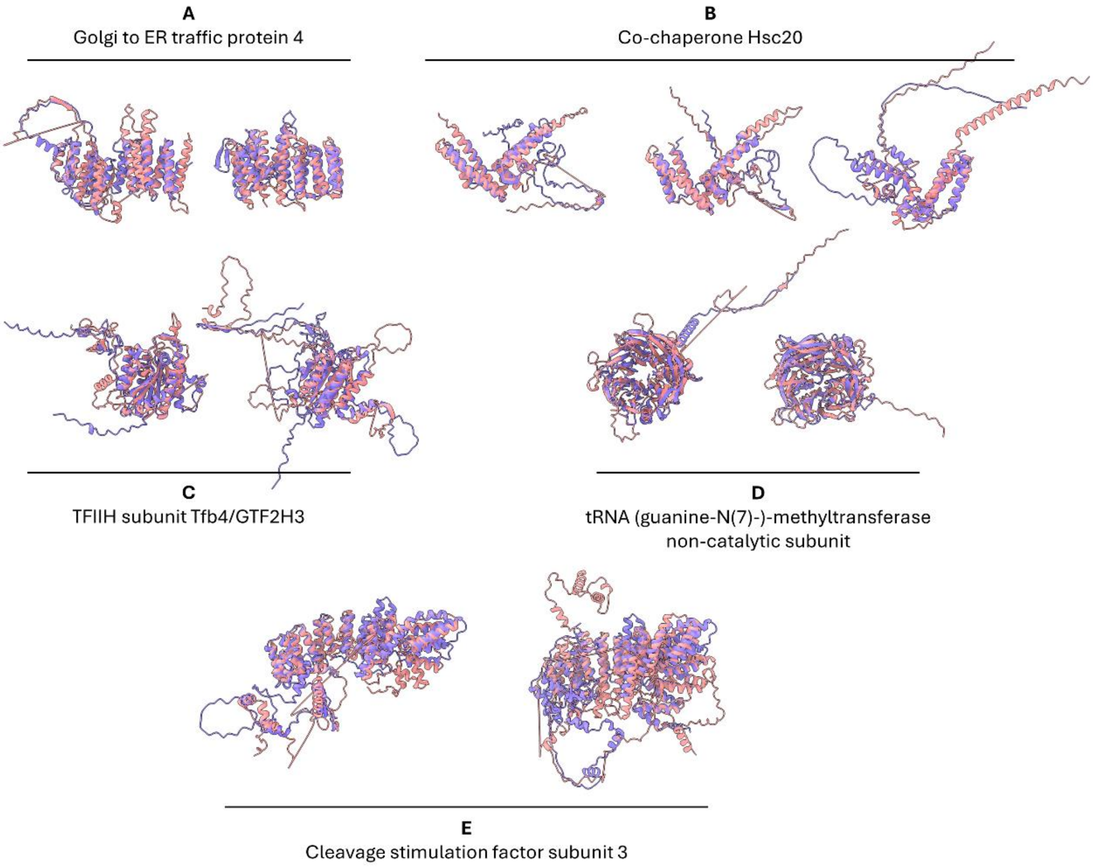


**Supplementary Figure 2. Essential eukaryotic proteins annotated by our approach that have been divided in subclusters by MMseqs2.** The figure shows 'BUSCO groups' that obtained more than one SRBH with the kinetoplastid proteins. The target protein (model species) is shown in violet, and the query protein (kinetoplastids) is shown in pink. In most cases, there are SRBH with homologs of this protein in different organisms we chose one as an example.

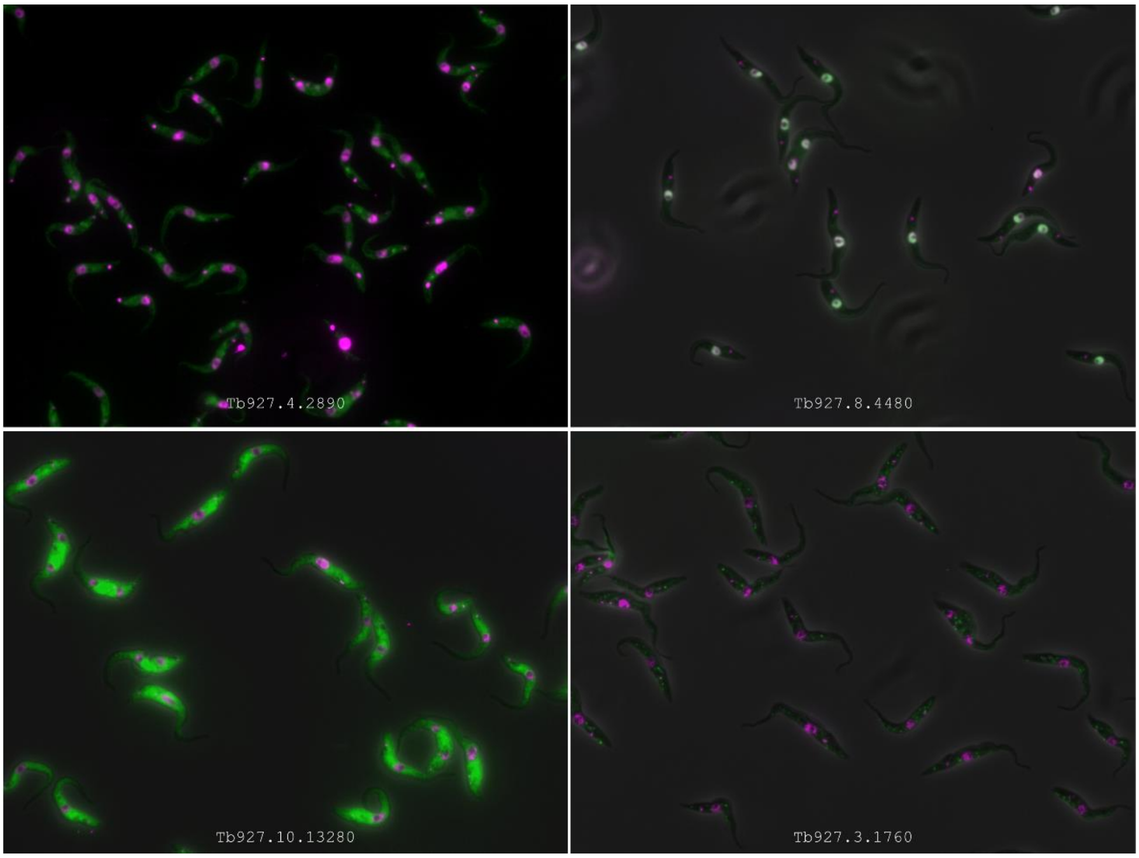


**Supplementary Figure 3. Subcellular Localization of selected *T. brucei* proteins in TrypTag.** Fluorescence microscopy images of *T. brucei* proteins (GFP tagged) from the clusters referred to as “missing gears,” showing their subcellular localization as determined by TrypTag. Upper panel: Left: Tb927.4.2890: Cytoplasm (weak; points), E3 ubiquitin-protein transferase MAEA (Q4D4T7) Right: Tb927.8.4480: Nucleoplasm, RNA polymerase II subunit A C-terminal domain phosphatase SSU72 (Q4DKS2). Lower panel: Left: Tb927.10.13280: Cytoplasm, eIF-2-alpha kinase activator GCN1 (A4HXV6). Right: Tb927.3.1760: Cytoplasm (points), Co-chaperone Hsc20 family protein (Q57ZD5).
