## Supplementary Tables for "Expanding kinetoplastid genome annotation through protein structure comparison"

**Supplementary Table 1.** Proteome description of model organisms used in this work.

| Species | Taxonomic lineage | Proteome Id | Protein count (UniProt) | Predicted structures (AFDBv4) | Genome assembly ID | BUSCO |
|---|---|---|---|---|---|---|
| **Methanocaldococcus jannaschii** | Archaea, Euryarchaeota, Methanomada group | UP000000805 | 1787 | 1773 | GCA_000091665.1 | C:98.2%S:98.0%,D:0.2%,F:0.4%,M:1.4%,n:958 |
| **Helicobacter pylori** | Bacteria, Campylobacterota, Epsilonproteobacteria | UP000000429 | 1554 | 1538 | GCA_000008525.1 | C:99.5%S:99.5%,D:0.0%,F:0.3%,M:0.2%,n:628 |
| **Campylobacter jejuni** | Bacteria, Campylobacterota, Epsilonproteobacteria | UP000000799 | 1623 | 1620 | GCA_000009085.1 | C:100.0%S:100.0%,D:0.0%,F:0.0%,M:0.0%,n:628 |
| **Neisseria gonorrhoeae** | Bacteria, Pseudomonadota, Betaproteobacteria | UP000000535 | 2106 | 2106 | GCA_000006845.1 | C:97.6%S:97.6%,D:0.0%,F:0.4%,M:2.0%,n:804 |
| **Klebsiella pneumoniae** | Bacteria, Pseudomonadota, Gammaproteobacteria | UP000007841 | 5728 | 5727 | GCA_000240185.2 | C:96.6%S:96.1%,D:0.5%,F:1.8%,M:1.6%,n:440 |
| **Haemophilus influenzae** | Bacteria, Pseudomonadota, Gammaproteobacteria | UP000000579 | 1704 | 1662 | GCA_000027305.1 | C:94.2%S:93.6%,D:0.5%,F:0.9%,M:4.9%,n:1100 |
| **Escherichia coli** | Bacteria, Pseudomonadota, Gammaproteobacteria | UP000000625 | 4404 | 4363 | GCA_000005845.2 | C:100.0%S:99.1%,D:0.9%,F:0.0%,M:0.0%,n:440 |
| **Shigella dysenteriae** | Bacteria, Pseudomonadota, Gammaproteobacteria | UP000002716 | 3897 | 3893 | GCA_000012005.1 | C:98.9%S:98.6%,D:0.2%,F:0.0%,M:1.1%,n:440 |
| **Salmonella typhimurium** | Bacteria, Pseudomonadota, Gammaproteobacteria | UP000001014 | 4533 | 4526 | GCA_000006945.2 | C:100.0%S:99.8%,D:0.2%,F:0.0%,M:0.0%,n:440 |
| **Pseudomonas aeruginosa** | Bacteria, Pseudomonadota, Gammaproteobacteria | UP000002438 | 5563 | 5556 | GCA_000006765.1 | C:99.5%S:99.0%,D:0.5%,F:0.3%,M:0.3%,n:782 |
| **Mycobacterium tuberculosis** | Bacteria, Terrabacteria group, Actinomycetota | UP000001584 | 3995 | 3988 | GCA_000195955.2 | C:99.1%S:98.5%,D:0.5%,F:0.4%,M:0.5%,n:743 |
| **Mycobacterium ulcerans** | Bacteria, Terrabacteria group, Actinomycetota | UP000020681 | 9033 | 9033 | GCA_000524035.1 | C:60.0%S:59.4%,D:0.7%,F:24.9%,M:15.1%,n:743 |
| **Nocardia brasiliensis** | Bacteria, Terrabacteria group, Actinomycetota | UP000006304 | 8414 | 8372 | GCA_000250675.3 | C:99.6%S:98.1%,D:1.5%,F:0.3%,M:0.1%,n:743 |

| | | | | | | |
|---|---|---|---|---|---|---|
| **Mycobacterium leprae** | Bacteria, Terrabacteria group, Actinomycetota | UP000000806 | 1603 | 1602 | GCA_000195855.1 | C:88.0%S:88.0%,D:0.0%,F:0.5%,M:11.4%,n:743 |
| **Enterococcus faecium** | Bacteria, Terrabacteria group, Bacillota | UP000325664 | 2823 | 2823 | GCA_008728475.1 | C:100.0%S:99.3%,D:0.7%,F:0.0%,M:0.0%,n:402 |
| **Staphylococcus aureus** | Bacteria, Terrabacteria group, Bacillota | UP000008816 | 2889 | 2888 | GCA_000013425.1 | C:98.4%S:98.4%,D:0.0%,F:1.3%,M:0.2%,n:450 |
| **Streptococcus pneumoniae** | Bacteria, Terrabacteria group, Bacillota | UP000000586 | 2031 | 2030 | GCA_000007045.1 | C:100.0%S:100.0%,D:0.0%,F:0.0%,M:0.0%,n:402 |
| **Dictyostelium discoideum** | Eukaryota, Amoebozoa, Evosea | UP000002195 | 12726 | 12622 | GCA_000004695.1 | C:93.7%S:89.8%,D:3.9%,F:1.6%,M:4.7%,n:255 |
| **Ajellomyces capsulatus** | Eukaryota, Opisthokonta, Fungi | UP000001631 | 9214 | 9199 | GCA_000150115.1 | C:96.4%S:95.7%,D:0.8%,F:1.7%,M:1.9%,n:4862 |
| **Paracoccidioides lutzii** | Eukaryota, Opisthokonta, Fungi | UP000002059 | 8811 | 8794 | GCA_000150705.2 | C:97.2%S:97.1%,D:0.1%,F:1.4%,M:1.4%,n:4862 |
| **Cladophialophora carrionii** | Eukaryota, Opisthokonta, Fungi | UP000094526 | 11181 | 11170 | GCA_001700775.1 | C:96.0%S:95.9%,D:0.1%,F:2.0%,M:2.0%,n:6265 |
| **Sporothrix schenckii** | Eukaryota, Opisthokonta, Fungi | UP000018087 | 8673 | 8652 | GCA_000474925.1 | C:95.8%S:95.6%,D:0.2%,F:1.1%,M:3.1%,n:3817 |
| **Fonsecaea pedrosoi** | Eukaryota, Opisthokonta, Fungi | UP000053029 | 12525 | 12509 | GCA_000835455.1 | C:99.3%S:99.1%,D:0.2%,F:0.3%,M:0.4%,n:6265 |
| **Candida albicans** | Eukaryota, Opisthokonta, Fungi | UP000000559 | 6036 | 5974 | GCA_000182965.3 | C:98.8%S:98.4%,D:0.5%,F:0.7%,M:0.5%,n:2137 |
| **Madurella mycetomatis** | Eukaryota, Opisthokonta, Fungi | UP000078237 | 9733 | 9561 | GCA_001275765.2 | C:89.3%S:79.5%,D:9.8%,F:0.9%,M:9.8%,n:3817 |
| **Saccharomyces cerevisiae** | Eukaryota, Opisthokonta, Fungi | UP000002311 | 6060 | 6039 | GCA_000146045.2 | C:99.6%S:97.4%,D:2.2%,F:0.1%,M:0.3%,n:2137 |
| **Schizosaccharomyces pombe** | Eukaryota, Opisthokonta, Fungi | UP000002485 | 5117 | 5128 | GCA_000002945.2 | C:81.8%S:79.0%,D:2.8%,F:1.4%,M:16.8%,n:1706 |
| **Onchocerca volvulus** | Eukaryota, Opisthokonta, Metazoa | UP000024404 | 12225 | 12047 | GCA_000499405.2 | C:98.2%S:84.0%,D:14.2%,F:0.5%,M:1.3%,n:3131 |
| **Schistosoma mansoni** | Eukaryota, Opisthokonta, Metazoa | UP000008854 | 10770 | 13865 | | C:78.7%S:64.9%,D:13.8%,F:2.6%,M:18.7%,n:954 |
| **Wuchereria bancrofti** | Eukaryota, Opisthokonta, Metazoa | UP000270924 | 13000 | 12721 | GCA_900622535.1 | C:77.8%S:77.3%,D:0.6%,F:7.1%,M:15.0%,n:3131 |
| **Dracunculus medinensis** | Eukaryota, Opisthokonta, Metazoa | UP000274756 | 10868 | 10834 | GCA_900625125.1 | C:78.1%S:77.5%,D:0.6%,F:5.8%,M:16.1%,n:3131 |
| **Brugia malayi** | Eukaryota, Opisthokonta, Metazoa | UP000006672 | 15168 | 8743 | | C:98.9%S:68.0%,D:30.9%,F:0.2%,M:0.9%,n:3131 |
| **Rattus norvegicus** | Eukaryota, Opisthokonta, Metazoa | UP000002494 | 47930 | 21270 | GCA_015227675.2 | C:98.1%S:45.6%,D:52.6%,F:0.3%,M:1.6%,n:13798 |
| **Mus musculus** | Eukaryota, Opisthokonta, Metazoa | UP000000589 | 54910 | 21615 | GCA_000001635.9 | C:99.8%S:51.3%,D:48.5%,F:0.0%,M:0.2%,n:13798 |
| **Homo sapiens** | Eukaryota, Opisthokonta, Metazoa | UP000005640 | 82493 | 23391 | GCA_000001405.29 | C:99.5%S:37.7%,D:61.8%,F:0.0%,M:0.5%,n:13780 |
| **Drosophila melanogaster** | Eukaryota, Opisthokonta, Metazoa | UP000000803 | 22049 | 13458 | GCA_000001215.4 | C:100.0%S:41.8%,D:58.2%,F:0.0%,M:0.0%,n:3285 |
| **Danio rerio** | Eukaryota, Opisthokonta, Metazoa | UP000000437 | 46578 | 24664 | GCF_000002035.6 | C:97.7%S:64.1%,D:33.6%,F:0.6%,M:1.7%,n:3640 |

| | | | | | | |
|---|---|---|---|---|---|---|
| **Caenorhabditis elegans** | Eukaryota, Opisthokonta, Metazoa | UP000001940 | 26708 | 19694 | GCA_000002985.3 | C:100.0%S:74.3%,D:25.7%,F:0.0%,M:0.0%,n:3131 |
| **Strongyloides stercoralis** | Eukaryota, Opisthokonta, Metazoa | UP000035681 | 12900 | 12613 | GCA_029582065.1 | C:69.9%S:67.1%,D:2.8%,F:0.8%,M:29.4%,n:3131 |
| **Plasmodium falciparum** | Eukaryota, Sar, Alveolata | UP000001450 | 5361 | 5187 | GCA_000002765.3 | C:99.1%S:98.0%,D:1.1%,F:0.0%,M:0.9%,n:3642 |
| **Zea mays** | Eukaryota, Viridiplantae, Streptophyta | UP000007305 | 63256 | 39299 | GCA_902167145.1 | C:96.6%S:43.2%,D:53.4%,F:0.8%,M:2.6%,n:4896 |
| **Oryza sativa** | Eukaryota, Viridiplantae, Streptophyta | UP000059680 | 48898 | 43649 | GCA_001433935.1 | C:84.4%S:78.1%,D:6.3%,F:5.0%,M:10.6%,n:4896 |
| **Glycine max** | Eukaryota, Viridiplantae, Streptophyta | UP000008827 | 74862 | 55799 | GCA_000004515.4 | C:99.2%S:25.5%,D:73.7%,F:0.2%,M:0.6%,n:5366 |
| **Arabidopsis thaliana** | Eukaryota, Viridiplantae, Streptophyta | UP000006548 | 39280 | 27434 | GCA_000001735.1 | C:100.0%S:64.3%,D:35.7%,F:0.0%,M:0.0%,n:4596 |
| **Trichuris trichiura** | - | UP000030665 | - | 9564 | - | - |

**Supplementary Table 2.** Gene IDs of kinetoplastid genes inside the 5 BUSCO essential protein clusters with single SRBH.

| Transcription initiation factor TFIID subunit 2 142542at2759 | RNA polymerase II subunit A 1304061at2759 | Ubiquinol-cytochrome c chaperone, CBP3 1428265at2759 | CTLH, C-terminal LisH motif 1087488at2759 | Translational activator Gcn1 160593at2759 |
|---|---|---|---|---|
| LSCM4_07687 | TcIL3000.A.H_000605100 | CFAC1_200019900 | DQ04_01261000 | TM35_000034270 |
| JKF63_07538 | TcIL3000_8_4250 | LtaP29.1460 | Tb427_040031300 | Tb427.10.13280 |
| LmxM.12.1250 | Tb1125.8.4480 | Tc_MARK_3098 | TEOVI_000465600 | Tb427_100139900 |
| LTRL590_120014700 | Tb927.8.4480 | ECC02_007338 | TevSTIB805.4.3010 | BCY84_18780 |
| LMJFC_120019600 | Tb427_080049700 | LSM04_000858 | Tbg972.4.2870 | C3747_393g7 |
| LdCL_120020400 | TevSTIB805.8.4620 | LPMP_291340 | Tb927.4.2890 | C4B63_4g201 |
| LARLEM1108_120014700 | Tbg972.8.4240 | TcCL_Unassigned05796 | Tb1125.4.2890 | TevSTIB805.10.13920 |
| LGELEM452_120015700 | TEOVI_000095600 | Tbg972.3.4250 | Tb427.04.2890 | Tc_MARK_4918 |
| LINF_120016700 | Tb427.08.4480 | LINF_290018900 | C3747_25g139 | LSM04_007840 |
| LtaP12.1060 | TcCL_ESM06005 | TcBrA4_0128780 | BCY84_19026 | TcYC6_0116000 |
| LmjF.12.1250 | TcBrA4_0019340 | CUR178_04195 | TcCL_ESM08092 | Tbg972.10.16050 |
| LTULEM423_120014800 | TRSC58_05615 | PCON_0043550 | TcG_01971 | TcCLB.506357.130 |
| LtaPh_1210600 | TcCLB.508989.70 | LbrM.29.1380 | TcCLB.504253.20 | ECC02_001088 |
| LMJLV39_120014700 | C3747_125g26 | DQ04_00341020 | C4B63_20g20 | DQ04_00161130 |
| LdBPK_120830.1 | TcCL_NonESM13541 | TEOVI_000029700 | TcYC6_0088390 | TcBrA4_0084050 |
| LAEL147_000172000 | TcG_02193 | LAMA_000114800 | TcBrA4_0122600 | Tb927.10.13280 |
| LAMA_000192700 | BCY84_05375 | LAEL147_000528700 | ECC02_007096 | TcG_03271 |
| LBRM2903_200076400 | C3747_66g56 | Tb1125.3.3890 | TcCL_NonESM02399 | Tb1125.10.13280 |
| LPMP_120930 | Tc_MARK_9824 | TRSC58_06831 | TCSYLVIO_003916 | TcIL3000.A.H_000837000 |
| LbrM.12.2.000930 | ECC02_003063 | TM35_000092450 | TCSYLVIO_003917 | TEOVI_000423300 |
| LbrM.12.0930 | TcCLB.509569.160 | BCY84_15100 | TcCLB.507735.60 | LtaP18.0830 |
| LPAL13_120014300 | C4B63_21g49 | JKF63_00542 | C3747_225g7 | LAMA_000281400 |
| LpyrH10_23_0150 | TCDM_03304 | TvY486_0303210 | Tc_MARK_2612 | LAEL147_000266100 |
| Lsey_0082_0120 | TcYC6_0064470 | BSAL_59245 | LSM04_001868 | LENLEM3045_180013900 |

| | | | | |
|---|---|---|---|---|
| LMARLEM2494_120014900 | TCSYLVIO_000479 | LpyrH10_08_1640 | TvY486_0402750 | LmxM.18.0820 |
| LSCM1_07901 | DQ04_03141080 | Baya_040_0230 | TM35_000014310 | CUR178_07044 |
| CFAC1_010017900 | TvY486_0803950 | LENLEM3045_290019700 | | LPAL13_180011100 |
| EMOLV88_120015600 | LSM04_006502 | TcG_02892 | | LDHU3_18.1070 |
| | TM35_000252160 | LGELEM452_290019500 | | LmjF.18.0820 |
| | | Tb927.3.3890 | | LTRL590_180013600 |
| | | TcCL_NonESM06154 | | LINF_180013400 |
| | | LMARLEM2494_290019200 | | LTULEM423_180013500 |
| | | C3747_13g293 | | LMJLV39_180013700 |
| | | TcIL3000_3_2500 | | LdBPK_180820.1 |
| | | LdCL_290019000 | | LbrM.18.2.000910 |
| | | TRSC58_02287 | | LSCM1_06728 |
| | | TCSYLVIO_004299 | | LSCM4_07084 |
| | | TcYC6_0109600 | | EMOLV88_180012900 |
| | | LtaPh_2914600 | | LGELEM452_180013000 |
| | | LMJFC_290020700 | | LMARLEM2494_180013700 |
| | | TevSTIB805.3.4110 | | LBRM2903_180014500 |
| | | LSCM4_03585 | | LMJFC_180014900 |
| | | LdBPK_291390.1 | | LtaPh_1808300 |
| | | LBRM2903_290020300 | | LbrM.18.0910 |
| | | LmjF.29.1300 | | LdCL_180013300 |
| | | LDHU3_29.1890 | | LARLEM1108_180013400 |
| | | LMJLV39_290019400 | | LdBPK.18.2.000820 |
| | | EMOLV88_360060800 | | LPMP_180820 |
| | | TcIL3000.A.H_000341700 | | JKF63_06568 |
| | | Tb427.03.3890 | | Lsey_0013_0410 |
| | | LSCM1_04426 | | LpyrH10_13_1080 |
| | | LTRL590_290020000 | | Baya_089_0240 |

| |
|---|
| C3747_47g274 |
| TcCLB.509999.80 |
| TCDM_04301 |
| LARLEM1108_290019500 |
| LdBPK.29.2.001390 |
| Tb427_030041700 |
| LTULEM423_290019200 |
| Lsey_0083_0040 |
| LPAL13_290017700 |
| C4B63_10g146 |
| LmxM.08_29.1300 |

**Supplementary Table 3.** Gene IDs of kinetoplastid genes inside the 5 BUSCO essential protein clusters with multiple SRBH.

| Golgi to ER traffic protein 4 1030907at2759 | Co-chaperone Hsc20 1129824at2759 | TFIIH subunit Tfb4/GTF2H3 1220881at2759 | tRNA (guanine-N(7)-)-methyltransferase non-catalytic subunit 937275at2759 | Tetratricopeptide-like helical domain superfamily 331411at2759 |
|---|---|---|---|---|
| LbrM.13.2.001300 | LSM04_005221 | TRSC58_02409 | Tc_MARK_3604 | LbrM.32.2.004190 |
| LPAL13_200009800 | TM35_000111840 | TCDM_05800 | C3747_40g223 | LAEL147_000663700 |
| LAEL147_000708700 | TEOVI_000541400 | TcBrA4_0035310 | BCY84_10939 | LMJLV39_320047400 |
| LmxM.33.0540 | Tb427.03.1760 | TcYC6_0050670 | TCSYLVIO_004859 | LTRL590_320047700 |
| LMJLV39_340011600 | Tb927.3.1760 | BCY84_08330 | TcCL_NonESM09808 | LdCL_320047000 |
| LAMA_000734700 | TevSTIB805.3.1770 | Tc_MARK_3393 | TcYC6_0124730 | LtaP32.4120 |
| LMARLEM2494_340011200 | Tb1125.3.1760 | TcCLB.509073.60 | TCDM_10622 | LMJFC_320053000 |
| LBRM2903_200011400 | Tbg972.3.1610 | TcCLB.508707.149 | TcG_06461 | LAMA_000694300 |
| LINF_340011000 | TRSC58_02379 | C3747_234g54 | BCY84_06378 | LmjF.32.3950 |
| LdBPK.34.2.000560 | TCSYLVIO_006355 | TcCL_NonESM05049 | TcBrA4_0015470 | LbrM.32.4190 |
| LDHU3_34.0950 | TcYC6_0119320 | ECC02_004028 | C4B63_84g58 | LGELEM452_320047800 |
| LtaP34.0600 | TcCLB.510091.50 | TcCL_ESM05086 | ECC02_001902 | JKF63_02758 |
| LGELEM452_340011000 | BCY84_03923 | C4B63_80g28 | TcCL_ESM11043 | LmxM.31.3950 |
| LSCM4_00818 | TcSYL_0019370 | C3747_68g50 | C3747_60g161 | CUR178_02533 |
| LdBPK_340560.1 | BCY84_13531 | TcG_04138 | TcCLB.509639.20 | LSCM4_03084 |
| LtaPh_3406000 | C3747_20g82 | TCSYLVIO_005377 | TcCLB.507711.120 | LARLEM1108_320047800 |
| LENLEM3045_340010900 | C3747_45g247 | CFAC1_190042600 | TRSC58_00287 | LINF_320046700 |
| LdCL_340011200 | TcBrA4_0082700 | LtaP32.3070 | DQ04_00121260 | LPMP_324080 |
| CUR178_01385 | TCDM_03945 | LbrM.32.2.003130 | TcIL3000.A.H_000817700 | LDHU3_32.5270 |
| LPMP_200550 | TcCL_NonESM05362 | LGELEM452_320036700 | TcIL3000_10_10140 | LMARLEM2494_320048200 |
| LMJFC_340011900 | ECC02_002253 | LBRM2903_320039200 | TcIL3000.A.H_000824600 | LSCM1_02053 |
| LARLEM1108_340011500 | Tc_MARK_5057 | LAMA_000684000 | Tb427_100119200 | LtaPh_3241200 |
| LbrM.13.1300 | TcCL_ESM04768 | LMARLEM2494_320036900 | Tb427.10.11210 | LTULEM423_320048400 |
| EMOLV88_340010500 | TcG_06056 | LbrM.32.3130 | TEOVI_000133900 | LPAL13_320048100 |

| | | | | |
|---|---|---|---|---|
| LmjF.34.0540 | TcCLB.510421.300 | CUR178_02424 | TevSTIB805.10.11780 | LENLEM3045_320047900 |
| LTULEM423_340011000 | C4B63_11g76 | LARLEM1108_320036700 | Tb1125.10.11210 | LdBPK_324100.1 |
| LTRL590_340011100 | DQ04_01111060 | LDHU3_32.3810 | Tbg972.10.13550 | LdBPK.32.2.004100 |
| LmjF.34.0540 | TvY486_0301062 | LSCM4_02974 | Tb927.10.11210 | EMOLV88_320043300 |
| CFAC1_290034100 | JKF63_04675 | LPAL13_320036700 | | Lsey_0261_0050 |
| JKF63_01649 | Lsey_0012_0480 | LINF_320036000 | | CFAC1_300048100 |
| Lsey_0144_0120 | CFAC1_230016200 | LmxM.31.2885 | | LpyrH10_02_4410 |
| LpyrH10_05_0810 | Baya_095_0230 | LTULEM423_320037300 | | C4B63_27g161 |
| C3747_57g63 | LmxM.25.1690 | LSCM1_01941 | | Tc_MARK_5200 |
| TcYC6_0126640 | LtaP25.1770 | LENLEM3045_320037200 | | TcG_09286 |
| TcBrA4_0017580 | LtaPh_2517700 | LmjF.32.2885 | | TcCLB.504005.50 |
| TcCLB.510187.140 | LAMA_000495100 | LPMP_323030 | | LSM04_008793 |
| TCDM_06586 | LBRM2903_250025800 | LtaPh_3230700 | | TcYC6_0035890 |
| BCY84_22873 | LdBPK.25.2.001760 | LAEL147_000652600 | | TcCL_ESM12074 |
| C4B63_81g55 | LbrM.25.2310 | LdBPK_323030.1 | | BCY84_01300 |
| Tc_MARK_6187 | LdBPK_251760.1 | LMJFC_320039400 | | TCSYLVIO_006462 |
| C3747_114g44 | LSCM1_05385 | LMJLV39_320036400 | | ECC02_006056 |
| TCSYLVIO_005008 | LARLEM1108_250023400 | LTRL590_320036700 | | C3747_107g90 |
| ECC02_002003 | LPMP_251760 | LdCL_320036300 | | TcBrA4_0109500 |
| | LPAL13_000047100 | LdBPK.32.2.003030 | | C3747_74g56 |
| | LTRL590_250024000 | JKF63_02652 | | TM35_000312400 |
| | LMJFC_250027500 | LpyrH10_02_3320 | | DQ04_03381070 |
| | LAEL147_000417800 | Lsey_0010_0360 | | TCDM_14112 |
| | LSCM4_04526 | | | Tbg972.11.13490 |
| | LDHU3_25.2200 | | | Tb927.11.12050 |
| | LENLEM3045_250024000 | | | Tb427tmp.01.3870 |
| | LMARLEM2494_250023400 | | | Tb427_110137300 |
| | EMOLV88_250022800 | | | Tb1125.11.12050 |

| | |
|---|---|
| LdCL_250023300 | TEOVI_000908500 |
| LMJLV39_250024300 | TcIL3000.11.12680 |
| LmjF.25.1690 | TevSTIB805.11_01.12430 |
| LTULEM423_250024100 | |
| LGELEM452_250023900 | |
| LINF_250023500 | |
| CUR178_05280 | |
| LbrM.25.2.002310 | |
| LpyrH10_10_2160 | |
| LPAL13_330036000 | |
| LMJFC_330040500 | |
| LMARLEM2494_330034800 | |
| LSCM1_02777 | |
| LBRM2903_330037600 | |
| LGELEM452_330036400 | |
| LINF_330036200 | |
| LmjF.33.2690 | |
| EMOLV88_330032700 | |
| LDHU3_33.3970 | |
| LARLEM1108_330033000 | |
| LdBPK.33.2.002830 | |
| LmxM.32.2690 | |
| LMJLV39_330038000 | |
| LPMP_332810 | |
| JKF63_02300 | |
| LENLEM3045_330035800 | |
| LtaP33.2920 | |
| LTULEM423_330036100 | |

| |
|---|
| LSCM4_02597 |
| LdBPK_332830.1 |
| LdCL_330035400 |
| LbrM.33.2970 |
| LAEL147_000695400 |
| CUR178_02045 |
| LbrM.33.2.002970 |
| LTRL590_330035600 |
| LAMA_000723200 |
| LtaPh_3329200 |
| CFAC1_210037700 |
| Lsey_0151_0010 |
| LpyrH10_03_4980 |

**Supplementary Table 4.** Multiple SRBH results detailed information.

| Query uniport accession (kinetoplastid) | Organism in clusters | Cluster representative | BUSCO ID of model organism hit | BUSCO Description | Model Organism | Protein names (model organism) | Target uniport accession (Model Organism) | GO description TrypTagg | Sequence identitiy of structural aligment |
|---|---|---|---|---|---|---|---|---|---|
| **A0A640KRB8** | Leishmania: 26, Leptomonas: 2, Crithidia: 1, Endotrypanum: 1, Porcisia: 1 | LbrM.13.2.001300 | 1030907at2759 | Golgi to ER traffic protein 4 | AJECG, ,PARBA, ,SPOS1 | DUF410 domain-containing protein | C0P182, C1GXR3, U7PSW4 | | 0.111, 0.112, 0.091 |
| **A0A2V2WCY5** | Trypanosoma: 10 | C3747_57g63 | 1030907at2759 | Golgi to ER traffic protein 4 | BRUMA,CANAL,DICDI,PLAF7 | BMA-CEE-1,Golgi to ER traffic protein 4, Golgi to ER traffic protein 4 homolog,Uncharacterized protein | A0A0K0K0X7,A0A1D8PND7,Q54TH4,Q8IL82 | | 0.114, 0.133, 0.124, 0.122 |
| **Q4D4T7** | Trypanosoma: 23 | DQ04_01261000 | 1087488at2759 | CTLH, C-terminal LisH motif | MOUSE,RAT,HUMAN | E3 ubiquitin-protein transferase MAEA (EC 2.3.2.27) (Erythroblast macrophage protein) (Macrophage erythroblast attacher), E3 ubiquitin-protein transferase MAEA (EC 2.3.2.27) (Macrophage erythroblast attacher), E3 ubiquitin-protein transferase MAEA (EC 2.3.2.27) (Cell proliferation-inducing gene 5 protein) (Erythroblast macrophage protein) (Human lung cancer oncogene 10 protein) (HLC-10) (Macrophage erythroblast attacher) (P44EMLP) | Q4VC33,Q5RKJ1,Q7L5Y9 | cytoplasm(weak; points) | 0.142, 0.139 |
| **Q57ZD5** | Trypanosoma: 25 | LSM04_005221 | 1129824at2759 | Co-chaperone Hsc20 | ORYSJ | Co-chaperone Hsc20 family protein, expressed (Os12g0456200 protein) (cDNA clone:J023020P09, full insert sequence) | Q2QRM4 | cytoplasm(points) | 0.168 |
| **Q4Q9S1** | Leishmania: 27, Leptomonas: 2, Blechomonas: 1, Crithidia: 1, Endotrypanum: 1, Porcisia: 1 | JKF63_04675 | 1129824at2759 | Co-chaperone Hsc20 | DROME,RAT,HUMAN,ARATH | Heat shock protein cognate 20 (MIP14027p), HscB mitochondrial iron-sulfur cluster co-chaperone, Iron-sulfur cluster co-chaperone protein HscB (DnaJ | A8JNT7,D3ZME7,Q8IWL3,Q8L7K4 | | 0.215, 0.212, 0.226, 0.142 |

| | | | | | | | | | |
|---|---|---|---|---|---|---|---|---|---|
| | | | | | | homolog subfamily C member 20) Cleaved into: Iron-sulfur cluster co-chaperone protein HscB, cytoplasmic (C-HSC20); Iron-sulfur cluster co-chaperone protein HscB, mitochondrial, Iron-sulfur cluster co-chaperone protein HscB homolog (AtHscB) | | | |
| **A0A640KWU3** | Leishmania: 27, Leptomonas: 2, Crithidia: 1, Endotrypanum: 1, Porcisia: 1 | LPAL13_330036000 | 1129824at2759 | Co-chaperone Hsc20 | AJECG,PARBA | DnaJ domain-containing protein,J-type co-chaperone JAC1 | C0NFZ6,C1H2E3 | | 0.128, 0.149 |
| **Q4E262** | Trypanosoma: 14 | TRSC58_02409 | 1220881at2759 | TFIIH subunit Tfb4/GTF2H3 | SOYBN,SCHPO,YEAST | General transcription and DNA repair factor IIH subunit TFB4 (RNA polymerase II transcription factor B subunit 4), General transcription and DNA repair factor IIH subunit tfb4 (TFIIH subunit tfb4) (RNA polymerase II transcription factor B subunit 4), General transcription and DNA repair factor IIH subunit TFB4 (TFIIH subunit TFB4) (RNA polymerase II transcription factor B 34 kDa subunit) (RNA polymerase II transcription factor B p34 subunit) (RNA polymerase II transcription factor B subunit 4) | A0A0R0JLP0,O74366,Q12004 | | 0.152, 0.15, 0.091 |
| **A0A0N1IME3** | Leishmania: 27, Leptomonas: 2, Crithidia: 1, Porcisia: 1 | CFAC1_190042600 | 1220881at2759 | TFIIH subunit Tfb4/GTF2H3 | AJECG | General transcription and DNA repair factor IIH subunit TFB4 (TFIIH subunit TFB4) (RNA polymerase II transcription factor B subunit 4) | C0NNU2 | | 0.071 |
| **Q4DKS2** | Trypanosoma: 27 | TcIL3000.A.H_000605100 | 1304061at2759 | RNA polymerase II subunit A | ARATH,DROME | RNA polymerase II subunit A C-terminal domain phosphatase SSU72 (CTD phosphatase SSU72) (EC 3.1.3.16) | A0A1P8AMK1,Q9VWE4 | nucleoplasm | 0.117, 0.105 |
| **E9AGI7** | Leishmania: 23, Leptomonas: 2, Crithidia: 1, | LSCM4_07687 | 142542at2759 | Transcription initiation factor TFIID subunit 2 | 9EURO2,9EURO1,CANAL,PARBA,RAT | Transcription initiation factor TFIID subunit 2, Transcription initiation factor TFIID subunit 2 | A0A0D2DTU3,A0A1C1CSD0,A0A1D8PQF6,C1H2G2,F1LNY6 | | 0.113, 0.124, 0.122, |

| | | | | | | | | | | |
|---|---|---|---|---|---|---|---|---|---|---|
| | Endotrypanum: 1, Porcisia: 1 | | | | | (Transcription initiation factor TFIID 150 kDa subunit) | | | | 0.115, 0.099 |
| **A4I4I7** | Trypanosoma: 26, Leishmania: 26, Leptomonas: 2, Bodo: 1, Blechomonas: 1, Crithidia: 1, Endotrypanum: 1, Porcisia: 1, Paratrypanosoma: 1 | CFAC1_200019900 | 1428265at2759 | Ubiquinol-cytochrome c chaperone, CBP3 | SOYBN,YEAST,CANAL,ORYSJ | Ubiquinol-cytochrome c chaperone domain-containing protein, Protein CBP3, mitochondrial Os07g0490300 protein (cDNA clone:J013149P03, full insert sequence) | I1MDZ1,P21560,Q5AC35,Q7XHR1 | | | 0.134, 0.126, 0.131, 0.123 |
| **A4HXV6** | Leishmania: 27, Trypanosoma: 20, Leptomonas: 2, Blechomonas: 1, Endotrypanum: 1, Porcisia: 1 | TM35_000034270 | 160593at2759 | Translational activator Gcn1 | SOYBN,DICDI | TOG domain-containing protein eIF-2-alpha kinase activator GCN1 (GCN1-like protein 1) (Translational activator gcn1) | I1JRU2,Q54WR2 | | cytoplasm | 0.11, 0.067 |
| **A4I8L3** | Leishmania: 26, Leptomonas: 2, Crithidia: 1, Endotrypanum: 1, Porcisia: 1 | LbrM.32.2.004190 | 331411at2759 | Tetratricopeptide-like helical domain superfamily | DANRE,SCHPO,HUMAN,CAEEL,MOUSE | Cleavage stimulation factor subunit 3 (Cleavage stimulation factor, 3 pre-RNA, subunit 3), mRNA 3-end-processing protein rna14" Cleavage stimulation factor subunit 3 (CF-1 77 kDa subunit) (Cleavage stimulation factor 77 kDa subunit) (CSTF 77 kDa subunit) (CstF-77) Suppressor of forked domain-containing protein | F1QIB2,O14233,Q12996,Q19866,Q99LI7 | | | 0.14, 0.097, 0.123, 0.146, 0.124 |
| **Q4CY43** | Trypanosoma: 23 | C4B63_27g161 | 331411at2759 | Tetratricopeptide-like helical domain superfamily | DRAME,WUCBA,SCHMA,RAT | Suf domain-containing protein, Suppressor of forked domain-containing protein, Putative cleavage stimulation factor, Cleavage stimulation factor subunit 3 | A0A158Q688,A0A3P7DR84,A0A3Q0KPI3,F1M4W7 | | | 0.135, 0.133, 0.139, 0.142 |
| **Q4DZR6** | Trypanosoma: 15 | Tc_MARK_3604 | 937275at2759 | tRNA (guanine-N(7)-)-methyltransferase non-catalytic subunit | DANRE,SCHPO,YEAST,MOUSE | tRNA (guanine-N(7)-)-methyltransferase non-catalytic subunit wdr4 (WD repeat-containing protein 4), tRNA (guanine-N(7)-)-methyltransferase non-catalytic subunit trm82 (Transfer RNA methyltransferase 82), tRNA (guanine-N(7)-)-methyltransferase non-catalytic subunit | A4IGH4,O74863,Q03774,Q9EP82 | | | 0.17, 0.156, 0.149, 0.139 |

| | | | | | | | | |
|---|---|---|---|---|---|---|---|---|
| | | | | | | TRM82 (Transfer RNA methyltransferase 82), tRNA (guanine-N(7)-)-methyltransferase non-catalytic subunit , WDR4 (Protein Wuho homolog) (mWH) (WD repeat-containing protein 4) | | |
| **G0UXW8** | Trypanosoma: 9 | TcIL3000.A.H_000817700 | 937275at2759 | tRNA (guanine-N(7)-)-methyltransferase non-catalytic subunit | CANAL | tRNA (guanine-N(7)-)-methyltransferase non-catalytic subunit TRM82 (Transfer RNA methyltransferase 82) | Q5AH60 | 0.137 |
